# Ablation of carotid body activity reverses diabesity by improving white and brown adipose tissue sympathetic innervation and metabolism

**DOI:** 10.1101/2023.08.04.552063

**Authors:** Bernardete F. Melo, Joana F. Sacramento, Julien Lavergne, Fátima O. Martins, Daniela Rosendo-Silva, Clara Panzolini, Gonçalo M. Melo, Cláudia S. Prego, Aidan Falvey, Elena Olea, Paulo Matafome, Asuncion Rocher, Jesus Prieto-Lloret, Miguel C. Correia, Phillipe Blancou, Silvia V. Conde

## Abstract

Finding novel pathological mechanisms that lead to innovative strategies to treat obesity and its associated illness are critically needed. The carotid bodies (CB) are metabolic sensors whose dysfunction contributes to insulin resistance and glucose intolerance development. Herein, we find that the ablation of CB activity, through resection of carotid sinus nerve (CSN) promotes weight loss and restores metabolic function in high fat (HF) rodents, by increasing WAT basal metabolism and by restoring WAT and BAT sympathetic activation. Moreover, we found that CSN resection rescues adipose tissue sympathetic/catecholamine resistance present in obesity states. Additionally, we found that the CB signals integrated in the paraventricular nucleus of the hypothalamus are decreased in obesity states and that CSN denervation in HF animals restore neuronal activity in this region. By inducing energy expenditure via the increase in WAT and BAT metabolism, CB modulation might be used as a therapeutic target for obesity and dysmetabolism.

**Highlights:**

- Ablation of carotid body (CB) activity promotes weight loss
- Carotid body modulates white and brown adipose tissue metabolism
- Ablation of CB activity rescues adipose tissue sympathetic resistance in obesity
- Carotid sinus nerve resection restored the altered neuronal activity induced by obesity in the paraventricular nucleus of the hypothalamus

## Introduction

Obesity, the epidemic of the XXI century, is a major public health concern, affecting one third of the worldwide population ^1^. This condition and its associated comorbidities - cardiovascular disease (CVD), type 2 diabetes (T2D), hypertension (HT), obstructive sleep apnea (OSA) and some cancers - contribute significantly to mortality worldwide, as it is estimated to be responsible for 1 in 5 deaths today ^2^.

Obesity management has been a challenge for years as the therapeutics available are often poorly efficient at reducing weight and/or require significant efforts in lifestyle modifications to be effective, which can be difficult to implement and, most of the times, only transiently successful. Thus, there is a compelling need for the discovery of novel mechanistic approaches and pharmacological targets that could lead to more effective and personalized therapeutic interventions for obesity.

Carotid bodies (CB) are peripheral chemoreceptors that respond to hypoxia by increasing chemosensory activity in the carotid sinus nerve (CSN), causing hyperventilation and activation of the sympathoadrenal system ^3^. Currently CBs are also considered metabolic sensors implicated in the regulation of peripheral insulin sensitivity, glucose homeostasis and lipid metabolism: 1) CB chemosensory activity is increased in prediabetes and type 2 diabetes animal models and overweight prediabetes patients ^3,4^; 2) abolishment of CB activity via resection or electrical modulation of the CSN prevented and reversed the metabolic alterations induced by hypercaloric diets in rats ^5^ and; 3) hyperbaric oxygen therapy, used to treat diabetic foot and that dramatically reduces CB activity ^6^, improves glucose homeostasis in T2D patients ^7^. Moreover, we showed that the restoration of metabolism induced by CSN-resection in hypercaloric animal models involves the improvement of insulin signaling and glucose uptake in the liver and adipose tissue (AT) being associated also with a decrease in fat deposition and weight gain induced by the diet ^5^. Therefore, our animal and human data strongly support a role for the CB in adipose tissue metabolism regulation and in the development of obesity and suggest that the modulation of CB activity can be a therapeutic strategy to treat overweight and obesity.

Additionally, it is consensual that CB controls sympathetic activity ^8–10^ and its overactivation has been experimentally shown to be at the origin of metabolic deregulation ^3,5,11,12^. On the other hand, AT sympathetic innervation is responsible for the modulation of lipolysis and thermogenesis via norepinephrine (NE) ^13^ and dopamine release ^14^. Therefore, in the search of the mechanistic pathways involved in CB modulation of obesity-related dysfunctions the circuit CB-AT-sympathetic nervous system surges with high relevance. In agreement, activation of CB chemoreceptors inhibits the elevated levels of BAT sympathetic nerve activity evoked by hypothermia ^15^.

Herein, we tested the innovative hypothesis that CB is a key player in neural sympathetic circuits controlling the white (WAT) and brown adipose tissue (BAT) metabolism. We also predict that CB/CSN blockade will have an anti-obesity effect that is associated with increased function of the adipose tissues and the recovery of sympathetic integration. Moreover, we investigated how CB signals to the central nervous system control AT physiology.

We demonstrated that the CB is a key intervenient in the neurocircuitry sympathetic nervous system-AT connection, introducing a completely new therapeutic target for obesity management.

## Results

### CSN resection decreases weight gain and adipose tissue deposition in obese rodents

Submitting rodents to hypercaloric diets promotes alterations on body weight, SNS activity, blood pressure and glucose metabolism, that are very similar with the human condition ^16^. Therefore, to investigate the effects of CB activity abolishment on weight gain and body fat mass, we used a diet-induced obesity model, by submitting Wistar rats and C75BL/6J mice to 10 and 12 weeks, respectively, of a lipid-rich diet (HF diet) (Fig. 1A). Both rats and mice submitted to hypercaloric diet exhibit a higher growth curve (Fig. 1B left and right panels, respectively) and increased weight gain (Fig. 1C) and total fat amount (Fig. 1D) than the normal chow (NC) animals, fed with a standard diet. The increase in fat amount was accompanied by an increase in all the WAT depots studied (Fig. 1E). Furthermore, and as previously demonstrated, HF diet promoted a dysmetabolic state, characterized by insulin resistance, glucose intolerance, dyslipidemia, hyperinsulinemia and increased c-peptide levels in rats and mice (Table S1). Additionally, and confirming previous data by Ribeiro et al. ^3^, obese rats showed increased respiratory responses to hypoxia (Fig. S1) without alterations in the responses to hypercapnia. To evaluate the link between the CB and its connection to the central nervous system in the development of obesity, we abolished CB activity, through CSN resection. CSN resection was confirmed by the abolishment of responses to hypoxia (Fig. S1). We found that bilateral CSN chronic resection decreased weight gain in the HF animals of both species (Fig. 1C). This effect was clearer when we plot only the weight gain after surgery and the week immediately after surgery was excluded (Fig. 1C). CSN resection decreased weight gain in both NC and HF diet animals. Note that obese rats and mice submitted to HF diet and to CSN resection decreased by 31% and 39% weight gain, respectively, in comparison with HF sham animals (Fig. 1C), respectively. This decrease in weight gain in HF animals was accompanied by a decrease in the total amount of fat (24% for rats and 26% for mice) (Fig. 1D) and by a decrease in all WAT depots (Fig. 1E). As expected, and previously described, CSN resection also reverses dysmetabolism, with an improvement of glucose tolerance and a reversion of insulin resistance, hyperinsulinemia and dyslipidemia (Table S1) not only in rats but also in mice, meaning that the beneficial effects of CSN on metabolism are not specie specific. Coincident with the increase in total WAT amount and depots, both rats and mice presented increased adipocytes perimeter in their perienteric adipose tissue (Fig. 1G), effects that were decreased by CSN resection in rats (55%) and mice (28%). HF diet also promoted alterations in the BAT in rats and mice, shown by an increase in adipocytes perimeter (Fig. 1H). Interestingly, CSN resection increased the mass of BAT in rats (53%) and mice (84%) (Fig. 1F), contributing to a decrease in adipocyte perimeter of 22% and 15%, respectively (Fig. 1H) suggesting that amelioration of metabolism in HF animals might include increased energy expenditure.

**Figure 1.**
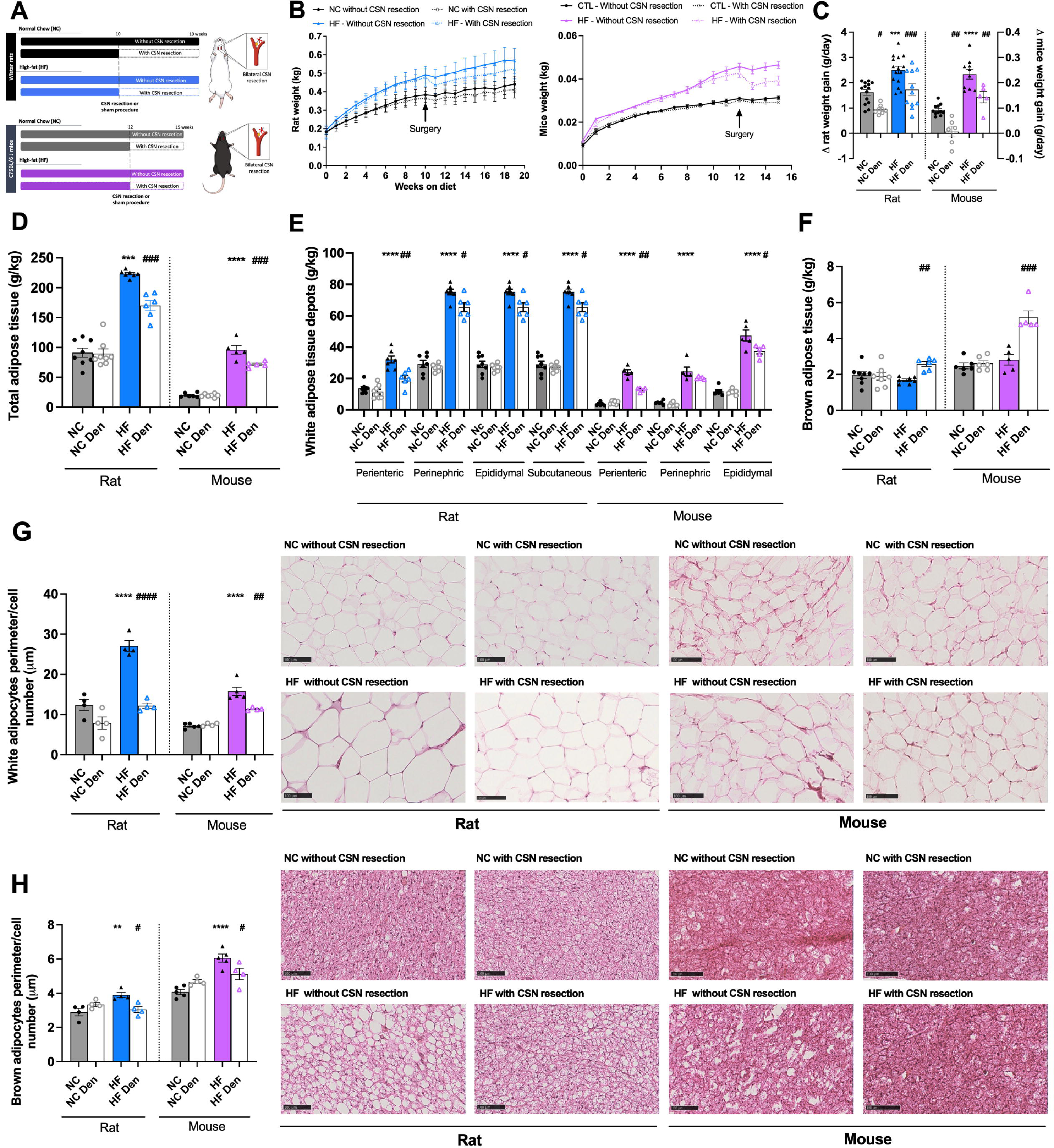
Carotid sinus nerve (CSN) resection decreases weight gain and adipose tissue deposition in obese rodents. A) Schematic representation of the protocol followed throughout the study for rats and mice. Effect of high fat (HF) diet and of CSN resection on: B) growth curves of Wistar rats (n=9-15, corresponding to two cohorts of animals that were used for different sets of experiments) (left panel) and C75BL/6 J mice (n=5-11) (right panel); C) average weight gain (g/day) in rats and mice after surgery; D) total white adipose tissue (WAT) weight (g/kg) (n=5-8); E) weight (g/kg) of white adipose tissue (WAT) depots in rats and mice (n=5-8); F) brown adipose tissue (BAT) weight (g/kg) (n=5-8); G) visceral WAT adipocytes perimeter per cell number (100 µm) - left panel shows average adipocytes perimeter, right panel shows representative H&E histological images of visceral fat in rat and mice (n=4-5); H) Brown adipocytes perimeter per cell number (100 µm) in the BAT depot in rats and mice (n=4-5) - left panel shows average adipocytes perimeter, right panel representative H&E histological images of BAT in rat and mice (n=4-5). Black and blue colors represent respectively, normal chow (NC) and HF diet rats. Grey and lavender colors show respectively NC and HF mice. Bars represent mean values ± SEM. Two-Way ANOVA with Bonferroni multicomparison test. \*\**p*<0.01, \*\*\**p*<0.001 and \*\*\*\**p*<0.0001 comparing NC vs HF groups; #*p*<0.05, ##*p*<0.01, ###*p*<0.001 and ####*p*<0.0001 comparing values with and without CSN resection.

### CSN resection ameliorates visceral WAT function in rodents by promoting the beiging of adipose tissue and an increase in its metabolism

We assessed visceral WAT metabolism by measuring basal oxygen consumption rate (OCR) and sympathetic-evoked OCR (NE [15µM] or dopamine [100nM]) using Seahorse technology in mice (Fig. 2A and 2B, left and right panels). It is well established that mitochondrial morphology, mass and function are impaired in multiple adipose tissue depots in obese rodents ^17^. In agreement, here we observed that the visceral WAT basal OCR of mice submitted to HF diet decreased by 51% in comparison with NC animals (Fig. 2C). Interestingly, CSN resection in HF animals restored WAT OCR by promoting an increase of 77% (Fig. 2C). Sympathetic mediators such as NE ^18,19^ or dopamine ^20,21^ activate WAT, and therefore we tested NE and DA on OCR and observed that NE and dopamine-evoked OCR were reduced by 53 and 44%, in WAT of HF mice in comparison with NC animals. CSN resection increased WAT responses to NE and dopamine both in NC and HF animals (55% in response to NE and 77% in response to dopamine) (Fig. 2D). To confirm a possible increase in adipose tissue energy expenditure mediated by CSN resection on obesity, we evaluated mitochondrial density and UCP1 expression (Fig. 2E) in the visceral WAT. HF diet decreased UCP1 levels (Fig. 2E, top panel) and mitochondrial density (Fig. 2E, bottom panel) in perienteric AT depot in rats and mice. Bilateral CSN resection increased UCP1 levels and mitochondrial density in both species, with an increase of 39 and 52% in UCP1, respectively, in obese rats and mice, and an increase in mitochondrial density of 61 and 52%, respectively, suggesting a restoration of WAT metabolism through increased thermogenesis. To confirm this phenotype, we assessed by western blot the levels of peroxisome proliferator activated receptor gamma coactivator 1 alpha (PGC1α) and peroxisome proliferator-activated receptor gamma (PPARγ) proteins, key mediators of lipid metabolism and markers of brown fat phenotype. HF diet decreased PGC1α levels by 27 and 67%, respectively, for rat and mice, effects completely reversed by CSN resection (Fig. 2G). Interestingly, HF diet did not alter the expression of PPARγ in visceral WAT of rats while decreased by 55% in the WAT of mice. CSN resection increased PPARγ expression by 73% in the HF rats and restored completely its expression in mice (Fig. 2H). Glucose uptake has been widely used as a surrogate marker for thermogenesis and energy balance, since glucose is one of the substrates for adipocytes metabolism ^20^. Here we evaluated glucose uptake, *in vivo*, in WAT depots, and the levels of hormone sensitive lipase (HSL) as well as the phosphorylated levels of adipose triglyceride lipase (ATGL) and AMP-activated protein kinase (AMPK) protein levels by Western Blot in rats (Fig. 3). HF diet decreased non-significantly glucose uptake in perienteric fat. CSN resection increased glucose uptake in all fat depots in normal chow and HF animals, an effect significantly different in the perienteric depot of the HF animals (Fig. 3B). Moreover, we found that HF diet decreased HSL without changing pATGL levels and that CSN increased HSL and pAGTL in both NC and HF animals (Fig. 3C and D), suggesting increased WAT lipid fluxes to generate FFA for thermogenesis (Fig. 3A). Finally, we found no alterations with HF diet in phosphorylated AMPK, that regulates both HSL and ATGL ^22^ of rats but a decrease of 34% in obese mice. CSN resection increased AMPK phosphorylation by 52 and 70%, respectively, in the CTL and HF Wistar rats, and restored the phosphorylation of AMPK in obese mice (Fig. 3E).

**Figure 2.**
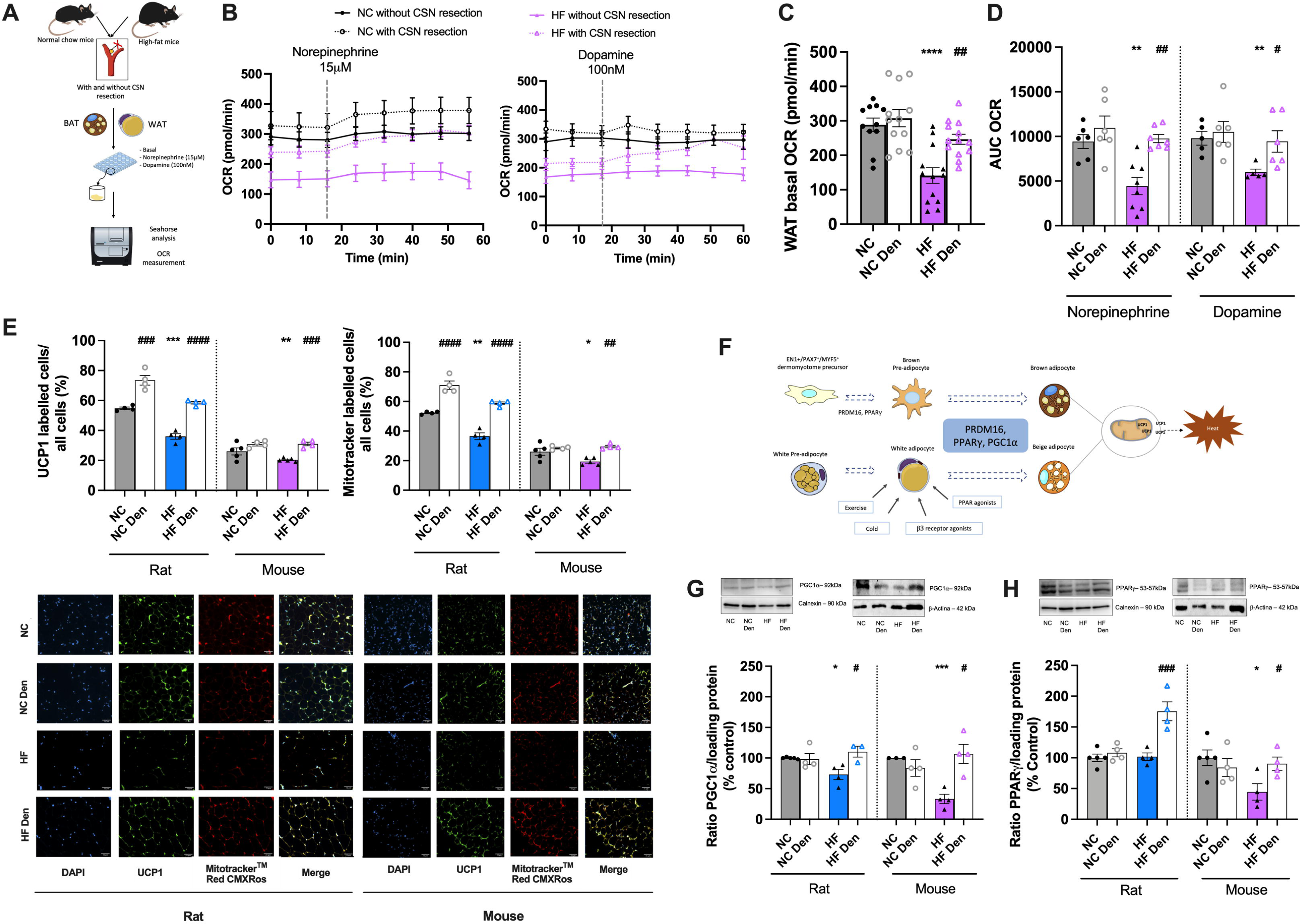
Carotid sinus nerve (CSN) resection ameliorates visceral white adipose tissue (WAT) metabolism in obese rodents. A) Schematic illustration of the protocol used to evaluate mice WAT and brown adipose tissue (BAT) oxygen consumption rate (OCR). Effect of high fat (HF) diet and of CSN resection on the: B) OCR per minute, before and after stimulation with norepinephrine [15µM] (left panel) or dopamine [100nM] (right panel) in mice (3 pieces of tissue from 4-6 animals); C) average basal OCR in mice (15-27 pieces of tissue from 4-6 animals); D) average OCR after stimulation with [15µM] or Dopamine [100nM] (3 pieces of tissue from 4-6 animals); E) percentage of UCP1 protein labeled cells and percentage of mitotrackerTM Red CMXRos labelled cells (top panels) in the perienteric depot (n=4-5) in rats and mice; bottom panels show representative images of UCP1 and MitotrackerTM Red CMXRos labelled cells, Green - UCP1 labelled adipocytes; Red - MitotrackerTM Red CMXRos labelled adipocytes; Blue - DAPI labelled nuclei of the adipocytes; Yellow - Merge of UCP1 and MitotrackerTM Red CMXRos labelled cells; F) Illustration of the molecular markers involved in brown adipocytes differentiation as well as the stimuli involved in the beiging of white adipose tissue; G) average expression of PGC1α (92kDa) (n=3-5) and H) average expression of PPARγ (53-57kDa) on visceral WAT of rats and mice (n=4-5) - representative western blots are shown on the top of the graphs. Grey and blue colors represent, respectively, normal chow (NC) and HF diet rats. Grey and lavender colors show NC and HF mice, respectively. Bars represent mean values ± SEM. Two-Way ANOVA with Bonferroni multicomparison test. \**p*<0.05, \*\**p*<0.01, \*\*\**p*<0.001 and \*\*\*\**p*<0.0001 comparing NC vs HF groups; #*p*<0.05, ##*p*<0.01, ###*p*<0.001 and ###*p*<0.0001 comparing values with and without CSN resection.

**Figure 3.**
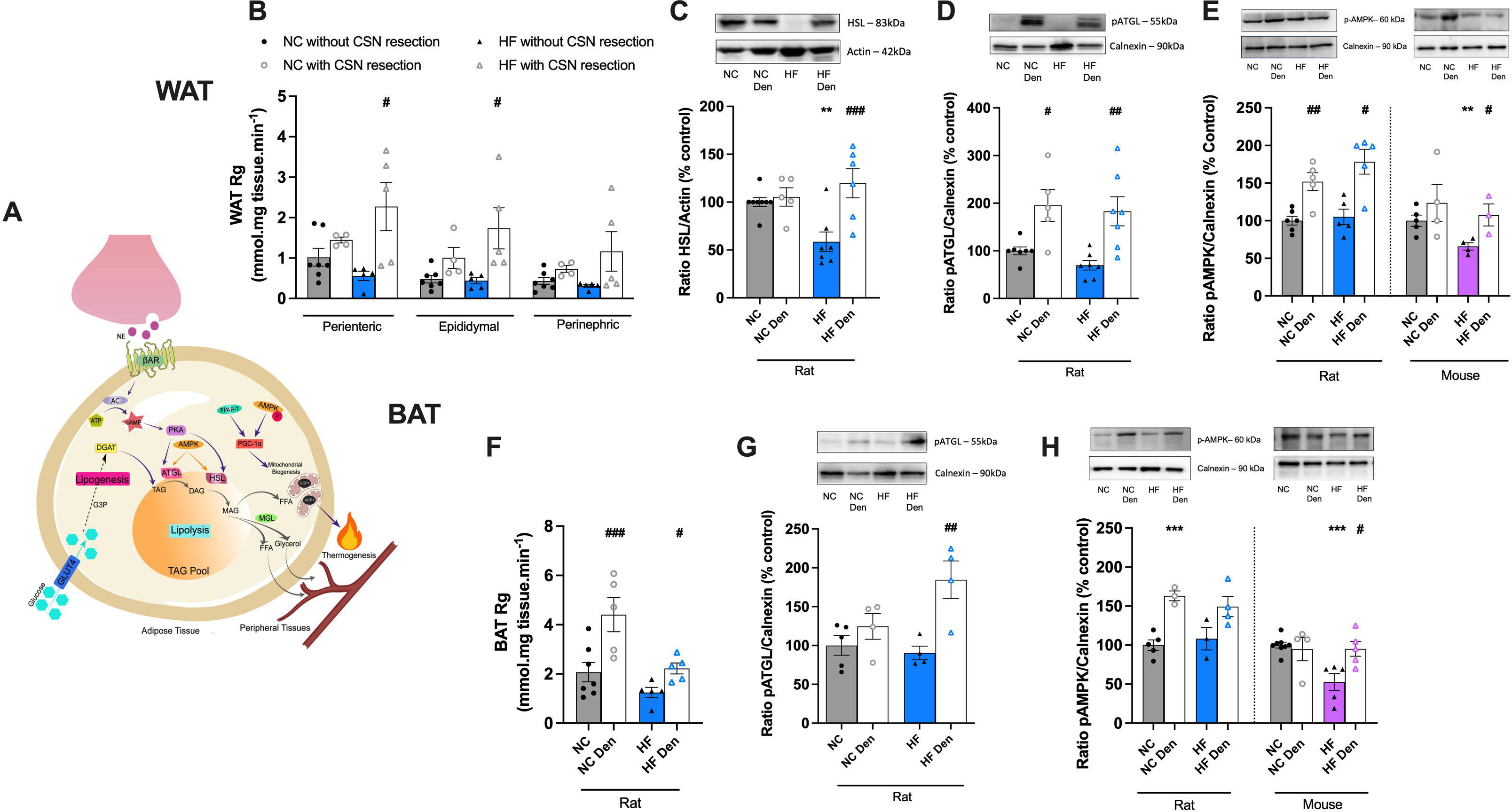
Carotid sinus nerve (CSN) resection increases lipid fluxes in white adipose tissue (WAT) and brown adipose tissue (BAT) in rodents. A) Illustration of the molecular markers involved in adipose tissue energy expenditure. Effect of high fat (HF) diet and of CSN resection on the: B) Rg’ values, reflecting glucose uptake on WAT depots in rats (n=4-7); C) average expression of HSL (83kDa) on WAT of rats (n=5-8); D) WAT average expression of phosphorylated ATGL (pATGL) (55kDa) in rats (n=5-7); E) WAT average expression of phosphorylated AMPK (pAMPK) (60kDa) in rats and mice (n=3-6) - representative western blots are shown on the top of the graphs; F) Rg’ values, reflecting glucose uptake on BAT depots in rats (n=5-7); G) BAT average expression of pATGL (55kDa) in rats (n=4-5); H) average expression of pAMPK (60kDa) on BAT of rats and mice (n=3-8) - representative western blots are shown on the top of the graphs. Grey and blue colors represent, respectively, normal chow (NC) and HF diet rats. Grey and lavender colors show NC and HF mice, respectively. Bars represent mean values ± SEM. Two-Way ANOVA with Bonferroni multicomparison test. \*\**p*<0.01 and \*\*\**p*<0.001 comparing NC vs HF groups; #*p*<0.05, ##*p*<0.01 and ###*p*<0.001 comparing values with and without CSN resection.

### CSN resection improves brown adipose tissue metabolism

It is consensual that obesity leads to BAT dysfunction with accumulation of enlarged lipid droplets and mitochondrial dysfunction ^23^. To study BAT mitochondrial function, we evaluated basal OCR as well as the OCR in response to NE [15µM] or dopamine [100nM] in mice (Fig. 4A, B and C). HF diet or with CSN resection did not change significantly basal OCR (Fig. 4A-B). As expected, NE increased OCR in BAT in NC animals (Fig. 4A, and C), an effect decreased by 20% in obese mice (Fig. 4C). CSN resection augmented NE activation of BAT by 23 and 42%, respectively in the NC and obese mice (Fig. 4C). Dopamine is described to directly activate thermogenesis and to increase mitochondrial mass in brown adipocytes ^24^. Herein, we were unable to see an increase in OCR evoked by dopamine both in NC and obese mice (Fig. 4A right panel). However, in CSN-resected animals’ dopamine increased OCR by 37% in mice fed with a standard diet, with no changes in obese mice (Fig. 4A-C). Looking for a correlation of OCR with increased thermogenesis, we evaluated mitochondrial density and the levels of proteins involved in the energy expenditure process by immunohistochenistry and Western blot. HF diet promoted a decrease of 21% in UCP1 labeled cells, in both species (Fig. 4D, top panel). In agreement, mitochondrial density decreased by 30 and 23%, respectively, in rats and mice submitted to the HF diet (Fig. 4D, bottom panel). Bilateral CSN resection increased UCP1 protein levels and mitochondrial density in NC animals and reversed the impact of HF diet in these parameters in both species. In contrast with what happened in visceral WAT, no alterations were observed in PGC1α (Fig. 4E) and PPARγ (Fig. 4F) protein levels with HF diet or CSN resection apart from the increase of 48% in PGC1α levels with CSN resection in mice (Fig. 4E). In agreement with a high metabolic activity, CSN-resected NC and obese animals exhibited an increased BAT glucose uptake, measured *in vivo* (Fig. 3F). Finally, and as it was observed in visceral WAT we found no alterations with HF diet in phosphorylated ATGL (Fig. 3G) and AMPK (Fig. 3H) in rats but a decrease of 48% in AMPK of obese mice. CSN resection increased the phosphorylation of ATGL by 104% (Fig. 3G) and AMPK by 38% in rats, and restored AMPK levels in mice, with an increase of 81% compared to the HF mice (Fig. 3H).

**Figure 4.**
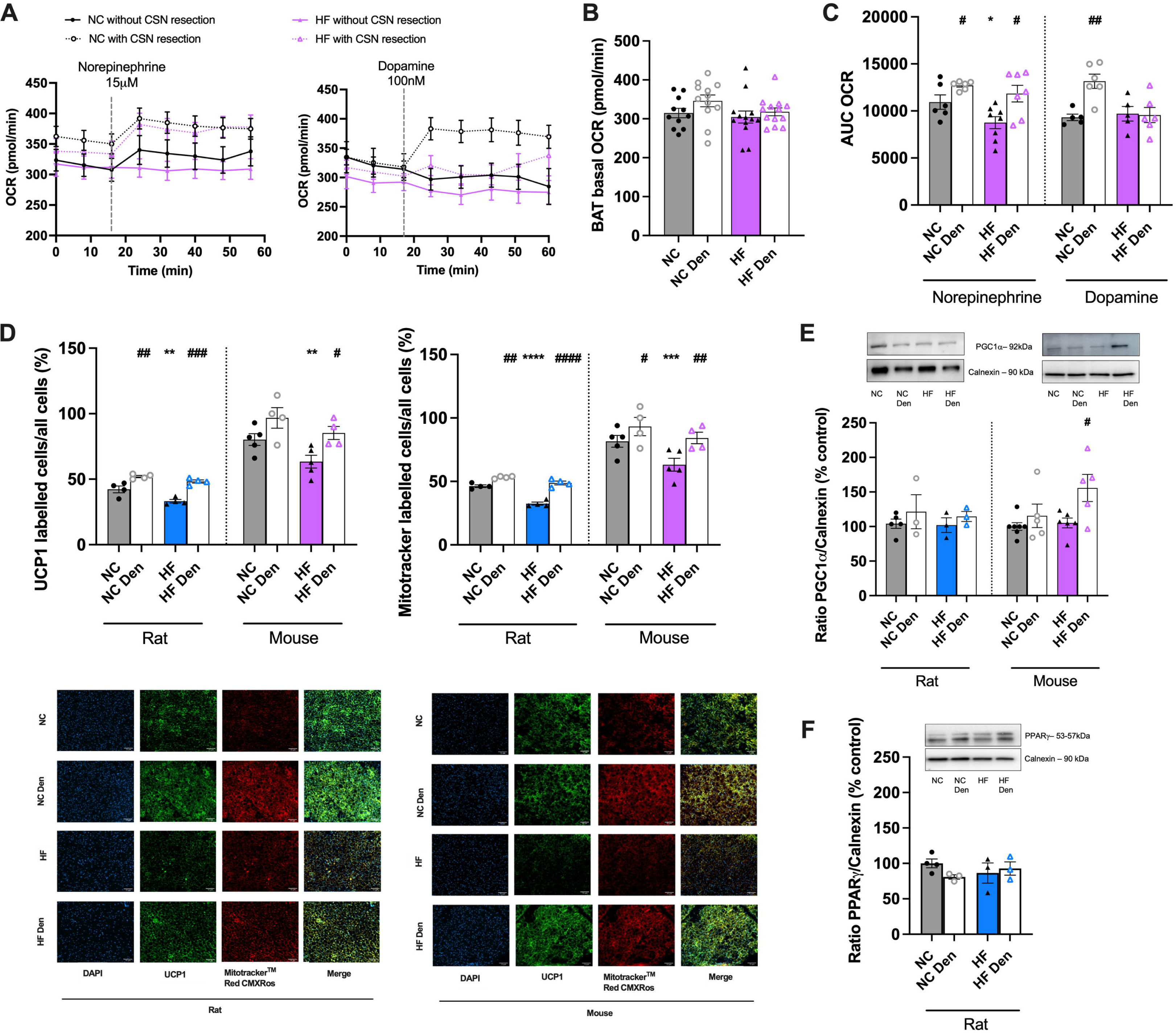
Carotid sinus nerve (CSN) resection improves brown adipose tissue (BAT) metabolism in rodents. Effect of high fat (HF) diet and of CSN resection on the: A) Curves of oxygen consumption rate (OCR) per minute, before and after stimulation with norepinephrine [15µM] (left panel) or dopamine [100nM] (right panel) in the BAT of mice (n=18-27 pieces of tissue from 6-8 animals). B) Average basal OCR in the BAT in mice (n=18-27 pieces of tissue from 6-8 animals); C) Average OCR after stimulation with norepinephrine [15µM] or dopamine [100nM] (n= 5-8 animals); D) percentage of UCP1 protein labeled cells and percentage of mitotrackerTM Red CMXRos labelled cells (top panels) in BAT (n=4-5) in rats and mice; bottom panels show representative images of UCP1 and MitotrackerTM Red CMXRos labelled cells, Green - UCP1 labelled adipocytes; Red - MitotrackerTM Red CMXRos labelled adipocytes; Blue - DAPI labelled nuclei of the adipocytes; Yellow - Merge of UCP1 and MitotrackerTM Red CMXRos labelled cells; E) average expression of PGC1α (92kDa) on BAT of rats and mice (n=3-7) and F) BAT average expression of PPARγ (53-57kDa) in rats (n=3-4) - representative western blots are shown on the top of the graphs. Grey and blue colors represent, respectively, normal chow (NC) and HF diet rats. Grey and lavender colors show NC and HF mice, respectively. Bars represent mean values ± SEM. Two-Way ANOVA with Bonferroni multicomparison test. \**p*<0.05, \*\**p*<0.01, \*\*\**p*<0.001 and \*\*\*\**p*<0.0001 comparing NC vs HF groups; #*p*<0.05, ##*p*<0.01, ###*p*<0.001 and ####*p*<0.0001 comparing values with and without CSN resection.

### CSN resection rescues adipose tissue sympathetic/catecholamine resistance in obesity

Obesity and dysmetabolic states have been linked to a whole-body overactivation of the SNS ^25^. In agreement, the obese rats exhibit an overactivation of SNS, reflected by an increase in 118 and 162%, respectively, in plasma NE and epinephrine levels (Table 1) and by an increase of 46% in the SNS index evaluated by heart rate variability analysis (Table 1). As expected, and in agreement with the findings that CB activation leads to an overactivation of the SNS ^12,26^, CSN resection normalized the effects of HF diet on plasma epinephrine and attenuated NE plasma levels and restored SNS index (Table 1). In contrast with the whole-body overactivation of the SNS (Table 1) and in line with findings that sympathetic activation increases AT metabolism ^27^, catecholamines levels within the visceral WAT, namely NE, epinephrine and dopamine were decreased by 55, 50 and 82%, respectively (Table 1). These effects were attenuated by CSN resection (Table 1). The decreased catecholamines content within the visceral WAT in obese animals were coincident with a decrease by 34% in the tyrosine hydroxylase (TH) levels in HF mice, evaluated by Western Blot (Fig. 5A and by a decrease of 83% in the intensity of TH innervation measured by light-sheet microscopy in the HF rats, with no significant alterations in the volume of fibers (Fig. 5B). Establishing a link between the CB and adipose tissue catecholaminergic signaling, CSN resection increased TH levels by 85% and 30% in obese rats and in mice, respectively (Fig. 5A) and increased by 174% the intensity of TH immunostaining, evaluated by light-sheetmicroscopy, without changing the volume of sympathetic fibers innervating visceral WAT (Fig. 5B, left panel). BAT followed the same tendency observed in visceral WAT, with the BAT of obese mice showing a decrease of 44% in TH levels (Fig. 5C), a decrease of 73% in the intensity of TH immunostaining in rat BAT and a decrease in the levels of NE (Table 1). Interestingly, HF diet induced a decrease of 73% in the volume of sympathetic fibers innervating the BAT (Fig. 5D). CSN resection in obese mice restored BAT TH levels to control levels (Fig. 5C) and attenuated the decrease in NE levels within the tissue (Table 1). In rats, CSN resection increased BAT TH expression (Fig. 5C) and the intensity of TH immunostaining, by 34% and 187%, respectively, in normal chow rats and TH immunostaining by 182% in HF animals, without changing the volume of sympathetic fibers innervating the BAT (Fig. 5D, left panel). These results indicate that the CB controls sympathetic innervation to the adipose tissue and that CSN resection restores catecholaminergic integration. Catecholamine resistance is a key feature of obese states ^28,29^ and is associated with altered levels of β-adrenergic receptors including the down-regulation of β3-adrenergic receptors ^29^. In agreement, we found that HF diet decreased β3 receptors levels by 13% (Fig. 5E, right panel) and promoted a decrease of both D1 and D2 receptors by 34 and 29%, respectively in WAT (Fig. 5F, left and right panels), while increasing β2 receptors by 26% (Fig. 5E, left panel). In BAT, HF did not change β2 levels while decreased β3, in rats, by 19%, while in mice the HF diet decrease β2 but increased β3 expression by 19% and 10%, respectively (Fig. 5G, left and right panels).

**Figure 5.**
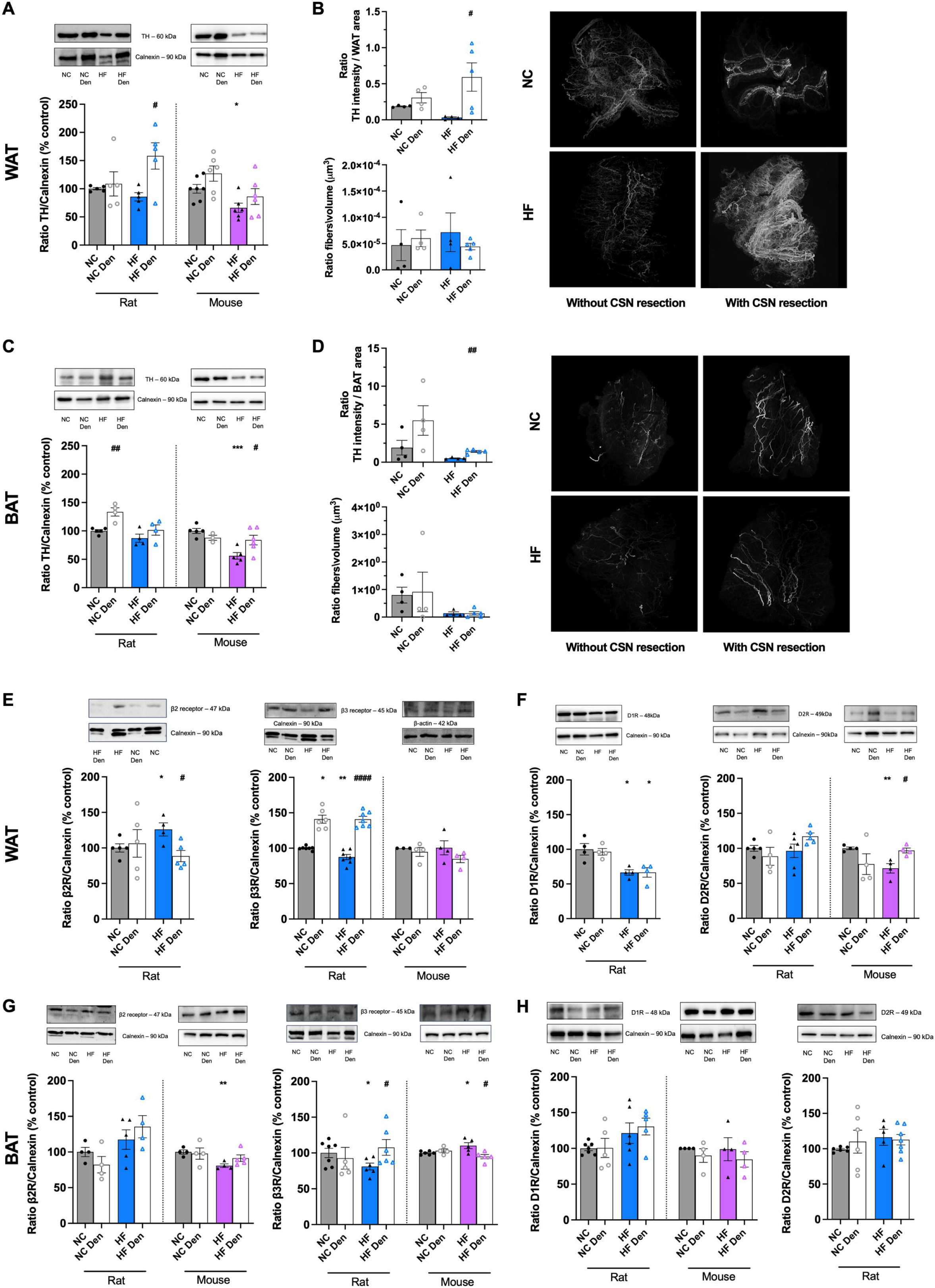
CSN resection restores sympathetic integration in visceral white (WAT) and brown (BAT) adipose tissues. Sympathetic innervation in the WAT and BAT presented by: A) WAT average TH expression (60 kDa) in rats and mice (n=5-7); B) intensity (left, top panel) and fibers volume (left, bottom panel) of TH immunolabeling (n=4-5) in WAT of rats – representative images are shown at the right panel, for animated gif of the images please consult https://fcmunlpt-my.sharepoint.com/:f:/g/personal/joana_sacramento_fcm_unl_pt/Ej7_uzyQyWBHqQVZQati9bkBFvDHgf1RTbGTEBihi4GzcQ?e=ta4OW2; C) BAT average TH expression (60 kDa) in rats and mice (n=4-6); D) intensity (left, top panel) and fibers volume (left, bottom panel) of TH immunolabeling (n=4-5) in BAT of rats – representative images are shown at the right panel; E) WAT average β2 receptors (□2R, 47 kDa) in rats (left panel) and β3 receptors (□3R, 45 kDa) in rats and mice (right panel) (n=3-7); F) WAT average dopamine type 1 receptors (□1R, 48 kDa) in rats (left panel) and dopamine type 2 receptors (D2R, 49 kDa) in rats and mice (left panel) (n=4-6); G) BAT average β2 receptors (□2R, 47k Da, left panel) and β3 receptors (□3R, 45 kDa, right panel) in rats and mice (n=3-7); H) average D1R (48 kDa) on the BAT of rats and mice (left panel) and D2R (49 kDa) on the BAT of rats (left panel) (n=4-7). Representative western blots are shown on the top of the graphs. Grey and blue colors represent, respectively, normal chow (NC) and HF diet rats. Grey and lavender colors show NC and HF mice, respectively. Bars represent mean values ± SEM. Two-Way ANOVA with Bonferroni multicomparison test. \**p*<0.05, \*\**p*<0.01 and \*\*\**p*<0.001 comparing NC vs HF groups; #*p*<0.05 and ##*p*<0.01 comparing values with and without CSN resection.

**Table 1.** Plasma and white and brown adipose tissue catecholamines levels in obese dysmetabolic rats.

| **Plasma Catecholamines**<br>**(pmol/ml)** | **Without CSN resection** | **With CSN resection** |
| --- | --- | --- |
| **Norepinephrine** |  |  |
| NC | 30.04±5.98 | 25.28±7.92 |
| HF | 65.52±15.83* | 50.16±10.23 |
| **Epinephrine** |  |  |
| NC | 41.17±5.97 | 46.23±11.94 |
| HF | 107.70±27.31* | 38.22±5.77 |
| **Dopamine + DOPAC** |  |  |
| NC | 1.93±0.18 | 2.30±0.67 |
| HF | 1.51±0.20 | 2.08±0.42 |
| **WAT Catecholamines**<br>**(pmol/mg tissue)** | **Without CSN resection** | **With CSN resection** |
| **Norepinephrine** |  |  |
| NC | 0.64 ± 0.04 | 0.68 ± 0.09 |
| HF | 0.29 ± 0.03**** | 0.52 ± 0.03<sup>#</sup> |
| **Epinephrine** |  |  |
| NC | 0.04 ± 0.01 | 0.05 ± 0.01 |
| HF | 0.02 ± 0.00** | 0.04 ± 0.01<sup>##</sup> |
| **Dopamine + DOPAC** |  |  |
| NC | 0.11 ± 0.02 | 0.07 ± 0.01 |
| HF | 0.02 ± 0.00**** | 0.07 ± 0.01<sup>###</sup> |
| **BAT Catecholamines**<br>**(pmol/mg tissue)** | **Without CSN resection** | **With CSN resection** |
| **Norepinephrine** |  |  |
| NC | 0.86±0.08 | 1.12±0.11 |
| HF | 0.53±0.10* | 0.73±0.13 |
| **Epinephrine** |  |  |
| NC | 0.04±0.02 | 0.03±0.01 |
| HF | 0.02±0.01 | 0.03±0.02 |
| **Dopamine + DOPAC** |  |  |
| NC | 0.02±0.00 | 0.03±0.01 |
| HF | 0.02±0.01 | 0.02±0.01 |
| **SNS Index** | **Without CSN resection** | **With CSN resection** |
| NC | 39.8±5.37 | 46.8±6.39 |
| HF | 58.0±5.22* | 38.2±5.57<sup>#</sup> |

Data are mean values ± SEM of 6 - 11 animals. Two-Way ANOVA with Bonferroni multicomparison test. \* $p<0.05$ , \*\* $p<0.01$ and \*\*\*\* $p<0.0001$ comparing CTL - normal chow - vs HF groups; # $p<0.05$ , ## $p<0.01$ and ### $p<0.001$ comparing values with and without CSN resection.

On the other hand, CSN resection in the WAT decreased β2 receptors expression by 30% while increased β3 expression by 61% in rats (Fig. 5D, left and right panel). In the BAT, CSN resection only altered β3 expression in the rats with an increase of 33% (Fig. 5G, right panel) and a decrease in the mice of 12% (Fig. 5G, right panel).

### CSN denervation restores neuronal activity in the paraventricular nucleus (PVN)

It is consensual that CB signals reaches the brain via integration in the nucleus tractus solitarius ^30^, and recently it was described that CB inputs are also integrated in the PVN ^31^. Therefore, we tested in rats if HF diet and CSN resection altered PVN activity, one of the most important autonomic control nuclei in the brain regulating metabolism and energy balance ^32^. We observed that HF diet promotes a decrease of 60% in the neuronal activity at -1.8mm coordinates at the PVN with no changes in the -2.16mm (Fig. 6B and 6C). Additionally, CSN resection in both coordinates of the PVN increased compared to control values the altered neuronal activity since CSN resection significantly increased by 131% (p<0.001) and 114% (p =0.06) the delta-FOS immunostaining in the -1.8mm and -2.16mm, respectively, in HF fed animals (Fig. 6A). Interestingly, CSN resection in NC animals at -2.16mm coordinates also increased by 102% (p= 0.09) neuronal activity.

**Figure 6.**
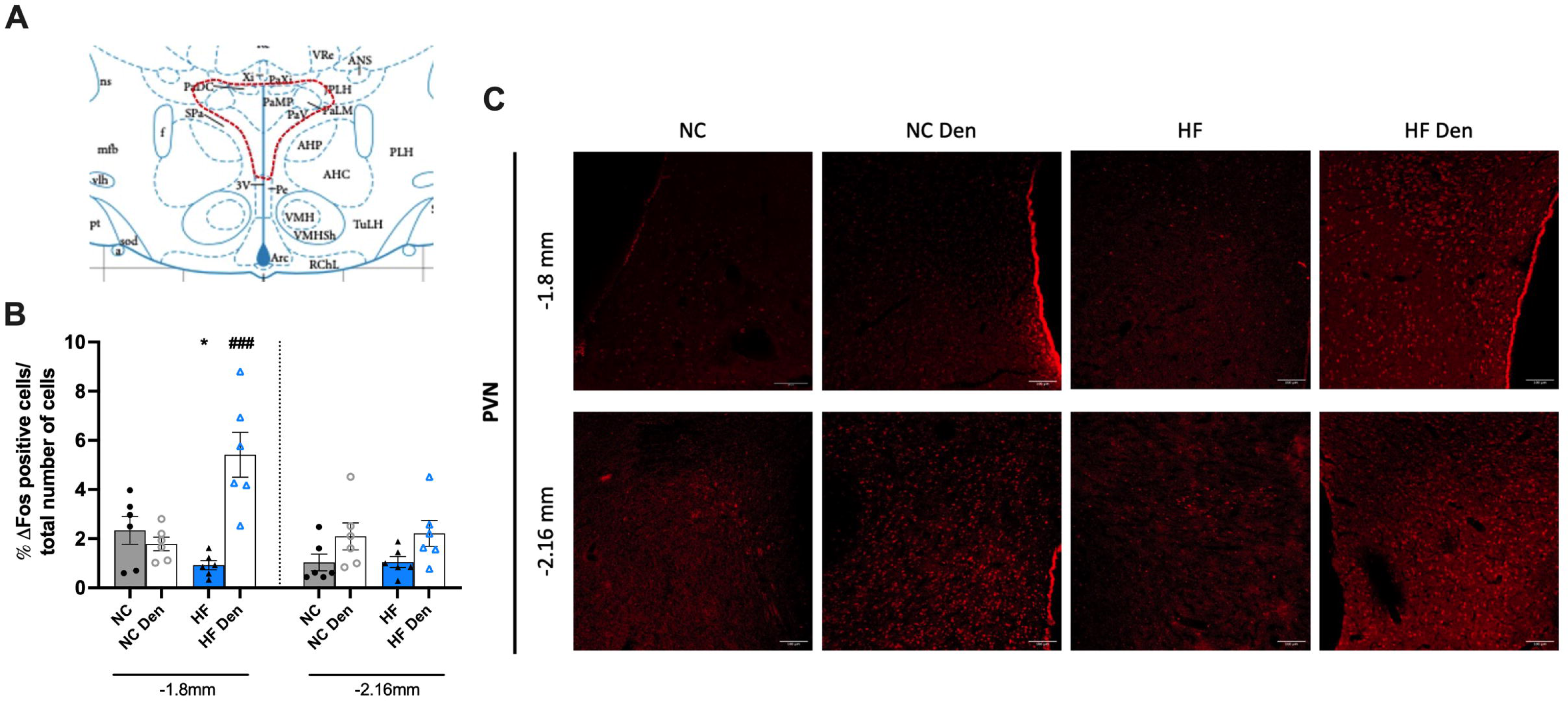
Ablation of carotid activity via CSN resection restores altered neuronal activity of hypothalamic paraventricular nucleus in obese dysmetabolic animals. Effect of high fat (HF) diet and of CSN resection on the: A) Illustration of the PVN region analyzed for the delta-FOS immunostaining obtained in the Paxinos & Watson Atlas for rat brain. B) Delta-FOS immunostaining in the PVN coordinates -1.8mm and -2.16mm from the bregma presented as percentage of delta-Fos positive cells/total number of cells (n=6) in rats. C) Representative images of immunostaining of delta-FOS for rats (scale = 100μm). Grey and blue colors represent, respectively, normal chow (NC) and HF diet rats. Bars represent mean values ± SEM. Two-Way ANOVA with Bonferroni multicomparison test. *p<0.05 comparing values with NC values; ###*p*<0.0001 comparing values with and without CSN resection.

## Discussion

Herein we described for the first time that the CB controls white and brown AT metabolism and restores sensitivity AT to catecholamines. Moreover, we show that this organ is a key player in the genesis and maintenance of obese states since the abolishment of its activity, via the resection of the CSN, decreases weight gain and adipose tissue deposition in obese rodents. CSN resection ameliorates baseline visceral WAT metabolism through the potentiation of energy expenditure, and via the improvement of BAT sympathetic activation. We demonstrate that CSN resection restores catecholaminergic action and sympathetic integration in visceral WAT and BAT in obese animals therefore concluding that the CB is a key intervenient in the neurocircuitry SNS-adipose tissue connection. Importantly, we also showed that all these mechanisms are well conserved in rodents. In agreement with the integration of CB inputs in the PVN, we show that CSN resection restored the altered neuronal activity induced by obesity in this nucleus of the hypothalamus, which is an important regulator of metabolism and energy expenditure.

### Effect of hypercaloric diets on weight gain and white and brown adipose tissue metabolism

The deleterious effects of hypercaloric diets are extensively described in the literature ^33,34^. Consistently, we showed that Wistar rats and C57BL/6 mice submitted to 19 or 15 weeks of 60% lipid-rich diet, respectively, presented a phenotype consistent with the phenotype observed in obese humans. They present a marked increase in weight gain accompanied by increased fat deposition, coincident with an increase in adipocytes perimeter both in WAT and BAT, and by the development of dysmetabolism characterized by insulin resistance, glucose intolerance, hyperinsulinemia, reduction on phosphorylated AMPK expression and dyslipidaemia. As expected, we show that 19 weeks of HF diet contributed to a decrease in both WAT and BAT metabolism, displayed by decreased basal OCR and OCR in response to catecholaminergic mediators as NE or dopamine, as well as a decrease in thermogenesis markers and a reduction on the expression of PGC1α and PPARγ. We also showed that in agreement with the increased weight gain and body fat mass in HF rodents, the increase in WAT depots was accompanied by increased adipocytes perimeter in the perienteric depot, as described in the literature in rodents ^35–37^ and in humans ^38^. Consistently with the metabolic dysfunction in HF animals, HF diet decreased BAT amount and contributed to increased lipid deposition within this tissue, with large lipid droplets being accumulated within brown adipocytes with increased adipocyte perimeter. The decrease in the BAT amount is not consensual with some studies using C57BL/6 mice and with different times of exposure to HF diet showing an increase in BAT weight ^39,40^. Nevertheless, the increase in adipocytes perimeter and the accumulation of lipid droplets within the brown adipocytes, that we observed, is consensual between several studies ^39,40^. These different results can be explained by differences in tissue collection and processing: in our work, the WAT around the BAT depot was dissected and weighed separately from the BAT depot, and in fact it was increased (data not shown), a detail that is not mentioned in these studies showing an increase in BAT weight.

Here we observed that WAT exhibits lower oxygen consumption rates than the BAT in lean animals, which is in line with the idea that adipose tissue oxygen consumption is relatively low in lean healthy subjects, accounting for approximately 5% of whole-body oxygen consumption ^41^. As expected, HF diet decreased WAT basal OCR, and reduced NE and dopamine-evoked OCR, suggesting that HF diet is not only affecting the mitochondrial activity of the adipocytes but also affecting adipocytes response to SNS stimuli and consequently affecting lipolysis ^41^. These results on the effects of HF diet on WAT metabolism are in accordance with the general idea that HF diet leads to a decreased WAT metabolism. In agreement, we found that UCP1 and mitotracker immuno-labeling, as well as PGC1α and PPARγ proteins were decreased in HF animals. Contrary, HF diet did not alter either BAT basal OCR or dopamine-stimulated OCR in obese mice but decreased the NE-evoked OCR in the BAT as well as UCP1 protein expression and mitochondrial density. In accordance with the decreased basal OCR activity in WAT and no effect on BAT induced by the HF diet, we also observed that HF diet decreased PGC1α, PPARγ and phosphorylated AMPK expression in the WAT, with no alterations in the BAT. In line with our results, some studies already demonstrated that in obese models, PGC1α mRNA levels were decreased not only in the WAT ^42^ and BAT ^43^ but also in skeletal muscle ^44^ and also that PPARγ is decreased in obese states ^45^.

### Mechanistic insights on the role of CB in obesity: linking the CB with catecholaminergic and metabolic activity of the adipose tissue

Obesity is associated with a general whole-body SNS overactivity, in particular in the outflow to the kidneys ^46^, with obese adults presenting increased urinary and plasmatic NA levels ^47^, and in the skeletal muscle vasculature ^48^. Moreover, animal ^49^ and clinical studies ^50^ demonstrated that increased SNS activity is involved in the pathophysiology of altered cardiac structure and function in essential hypertension, and that obese insulin-resistant individuals display blunted sympathetic neuronal responses to physiological hyperinsulinemia, glucose consumption and changes in energy states ^51^. In agreement with these studies and as shown by our group in animal models of prediabetes and T2D ^3,5^, we observed increased plasma levels of NE and epinephrine and whole-body SNS index in the obese rats, demonstrating overall SNS overactivity.

It is consensual that the SNS directly innervates both BAT and WAT and plays a key role in the modulation of lipolysis and/or energy expenditure/thermogenesis through NE and dopamine release ^14^ and the activation of its receptors ^13,52^. For example, β2 and β3 agonists have been shown to induce thermogenesis in the BAT ^53,54^ and dopamine have been shown to directly increase mitochondrial mass and thermogenesis in this tissue ^24^. More recently, dopamine was also shown to potentiate insulin action on glucose uptake in visceral WAT and to act to promote adipose tissue metabolism ^21^. Here we show that hypercaloric diet intake produce a decrease in baseline metabolism in WAT, assessed by measuring OCR, as well as a decrease in mitochondrial activity, UCP1 expression, PPARγ and PGC1α and decreased catecholaminergic activation in WAT and BAT. These results are in agreement with the decreased levels of catecholamines found in WAT and BAT and with the low intensity of TH expression in the adipose tissue measured by light-sheet microscopy. Despite these results contradict the idea that obesity and its metabolic comorbidities are associated with a SNS overactivation, recently several pieces of evidence were generated supporting the idea of a regional activation/modulation of the SNS ^55^. Our results clearly support that HF diet produced a decreased catecholaminergic activation to the adipose tissues while increasing whole-body SNS activity.

The CB is a powerful modulator of SNS activity, as shown by previous studies on its involvement in cardiovascular responses/adjustments to hypoxia ^9^ and its involvement on the pathophysiological mechanisms of sympathetic mediated diseases, as essential hypertension ^8,56^ and chronic heart failure ^57^. Moreover, the dysfunction of CB activity leading to the overactivation of the SNS has already been described by our group to be associated with the development and maintenance of prediabetes, T2D and metabolic syndrome ^3,5,58^. In fact, CSN resection, that abolish the connection between the CB and the central nervous system, prevents and normalizes sympathetic overaction measured as: plasma and adrenal medulla catecholamines content, the increase percentage of low frequencies, the ratio low frequency/high frequency in the power spectra of the heart rate variability ^59^ and the increase in the electrophysiological activity measured in the cervical sympathetic chain ^60^. Here we described that the CB is also a powerful modulator of catecholaminergic action/sympathetic integration in the adipose tissue, controlling adipose tissue metabolism, both white and brown. We showed that the abolishment of CB connectivity to the central nervous system, by resecting the CSN, decreases weight gain and adipose tissue deposition in obese rodents, contributing to a reduction of adipocytes perimeter and reversing metabolic dysfunction. Moreover, CSN resection ameliorated visceral WAT and BAT metabolism, increasing thermogenic markers and improving glucose metabolism, which is in line with the previous findings that CSN resection improved dysmetabolism by positively impacting glucose metabolism in visceral adipose tissue ^5^. The link between the BAT and CB chemoreceptors in not new, as Madden and Morrison ^15^ previously showed that activation of CB chemoreceptors inhibits the elevated levels of BAT sympathetic nerve activity evoked by hypothermia. The novelty highlighted in this paper is this link between the improvement of metabolism and decrease in weight gain by restoring WAT sensitivity to catecholamines and increasing BAT metabolism. Novelty also lies in the link between the CB and the SNS in the control of the WAT metabolism.

Importantly, we also show herein that the link between CB and BAT and WAT metabolism seems to involve the integration of CB inputs into the PVN of the hypothalamus as we described that CSN denervation augments the decreased neuronal activity in the PVN in obesity. The decreased neuronal activity in the PVN in dysmetabolic states is not new ^61^ and has been associated with increased inflammation ^62,63^, and with the loss of mitochondria and MC4R protein loss in Sim1/MC4R neurons ^64^. As well, CB inputs are known to be integrated in this nucleus ^31^ and melanocortin-induced activation of PVN was shown to affect sympathetic outflow to interscapular BAT and BAT thermogenesis ^65^, therefore it is plausible to expect this link between the CB, the PVN and BAT and possible with the WAT herein observed. Nevertheless, it remains to be established which neuronal populations within the PVN are being activated by CSN denervation in obesity and to completely decipher the neuronal circuits/projections integrating all the CB metabolic functions.

In conclusion this work adds to the understanding of CB role in the control of whole-body metabolism, its role in the genesis of metabolic dysfunction and the pathophysiological mechanisms behind it. Herein we tested the innovative hypothesis that the CB is involved in the genesis and maintenance of obesity states and that the abolishment of its activity can decrease weight gain and ameliorate obesity-comorbidities. Also, we show for the first time that the CB controls the catecholaminergic system and sympathetic integration on the white and brown adipose tissue depots, and therefore the targeting of CB and modulation of its activity have high potential as a therapeutic approach for the management of overweight and obesity states.

## Methods

### Diets and Animal Care

Experiments were performed in 8–9-weeks-old male Wistar rats (200–300 g) obtained from the animal house of NOVA Medical School, Faculty of Medical Sciences, Universidade NOVA de Lisboa, Portugal and 4 week-old male C75BL/6 J mice purchased from Charles River (Massachusetts, EUA). After randomization, the animals were assigned to the HF group, fed with 60% fat diet (61.6% fat + 20.3% carbohydrate + 19.1% protein; Test Diets, Missouri, USA) for 10 weeks (Wistar rats) or 12 weeks (C75BL/6 J mice); or to an aged-matched control group fed with a standard diet (7.4% lipid and 75% carbohydrates, of which 4% were sugars and 17% protein). After 10 and 12 weeks of diet for rats and mice, respectively, both HF and control groups were randomly divided, and half of the group was submitted to carotid sinus nerve (CSN) resection and the other half submitted to sham procedure, in which the CSN is left intact. These procedures were performed as previously described by Sacramento et al. ^5^.

After CSN resection, the groups were maintained under the respective diets followed for 9 weeks or 3 weeks, for rats and mice respectively. At a terminal experiment the animals were anesthetized with sodium pentobarbitone (60 mg/kg, i.p), and in some cohort’s catheters were placed in the femoral artery for arterial blood pressure measurement. Blood was collected by heart puncture for serum and plasma quantification of mediators (such as insulin and c-peptide levels and inflammatory markers) and the adipose tissue pads (visceral, perinephric, epididymal, subcutaneous depots and interscapular brown adipose tissue depot) were then rapidly collected, weighted, and stored at either –80 °C or transferred to 4% PFA or collected and maintained in Tyrode medium (140 mM NaCl, 5 mM KCl, 2 mM CaCl2, 1.1 mM MgCl2, 10 mM Hepes and 5.5 mM glucose; pH 7.4) on ice for O_2_ consumption rate analysis.

During all the experimental period animals were kept under temperature and humidity control (21D±D1 °C; 55 ± 10% humidity) with a 12 h light/12 h dark cycle and were given ad libitum access to food and water, except in the night prior to insulin sensitivity and glucose tolerance evaluation. Body weight was monitored weekly, and energy and liquid intake were monitored daily. Insulin sensitivity and glucose tolerance were also evaluated over the experimental period. CSN resection was confirmed by the abolishment of responses to hypoxia in conscious free-movement animals by pletismography.

Laboratory care was in accordance with the European Union Directive for Protection of Vertebrates Used for Experimental and Other Scientific Ends (2010/63/EU). Experimental protocols were approved by the NOVA Medical School/Faculdade de Ciências Médicas Ethics Committee (n°08/2015/CEFCM and 41/2021/CEFCM), by Portuguese Direção-Geral de Alimentação e Veterinária (DGAV) and by the National Ethical Committee of France (CIEPAL #2018082016528754).

### Insulin sensitivity and glucose tolerance evaluation

Insulin sensitivity was evaluated using an insulin tolerance test (ITT) after overnight fasting (Wistar rats) or 5 hours fasting (C75BL/6 J mice) as previously described ^5^. Glucose tolerance was evaluated using oral glucose tolerance test (OGTT) after overnight fasting followed by administration of a saline solution of glucose (2 g/kg or 1.5g/kg, for rats or mice, respectively; VWR Chemicals, Leuven, Belgium), by gavage, as described ^5^.

### Pletismography

Ventilation was measured in conscious freely moving rats as previously described by Sacramento et al. ^59^. The system (emka Technologies, Paris, France) consisted of 5-L methacrylate chambers continuously fluxed (2 L/min) with gases. Tidal volume (VT; ml), respiratory frequency (f; breaths/min; bpm), the product of these two variables, and minute ventilation (VE; ml/min/Kg) were monitored. Each rat was placed in the plethysmography chamber and allowed to breathe room air for 30 min until they adapted to the chamber environment and acquired a standard resting behavior. The protocol consisted in animal acclimatization during 30 min followed by 10 min of normoxia (20% O2 balanced N2), 10 min of hypoxia (10% O2 balanced N2), 10 min of normoxia, 10 min of hypercapnia (20% O2 + 5% CO2 balanced N2), and finally to 10 min of normoxia. A high-gain differential pressure transducer allows measuring the pressure change within the chamber reflecting tidal volume (VT). False pressure fluctuations due to rat movements were rejected. Pressure signals were fed to a computer for visualization and storage for later analysis with EMKA software (emka Technologies, Paris, France).

### Evaluation of Autonomic Nervous System

The balance between the sympathetic and parasympathetic components of the autonomic nervous system was made by calculating the sympathetic nervous system (SNS) and parasympathetic nervous system (PNS) indexes computed in Kubios HRV software (www.kubios.com). The SNS index in Kubios is based on Mean heart rate, Baevsky’s stress index, and low frequency power expressed in normalized units and the PNS index which is based on the mean intervals between successive heartbeats (RR intervals), the root mean square of successive RR interval differences (RMSSD) and high frequency power expressed in normalized units. Heart rate and RR intervals were obtained using Iox 2.9.5.73 software (Emka Technologies, Paris, France), with an acquisition frequency of 500 Hz.

### Quantification of Biomarkers: Plasma Insulin, C-Peptide, Lipid Profile, and Catecholamines

Insulin and C-peptide concentrations were determined with an enzyme-linked immunosorbent assay kit (Mercodia Ultrasensitive Rat Insulin ELISA Kit and Mercodia Rat C-peptide ELISA Kit, respectively, Mercodia AB, Uppsala, Sweden). Catecholamines were measured in plasma and in homogenized perienteric adipose tissue samples by HPLC with electrochemical detection as previously described ^66^ while plasma IL-10 and TNFα levels were evaluated using the chemiluminescence-based assay kit V-PLEX Proinflammatory Panel 1 Mouse Kit (Meso, Maryland, USA). Lipid profile was assessed using a RANDOX kit (RANDOX, Irlandox, Porto, Portugal).

### Western Blot Analysis

Visceral WAT (100 mg) and BAT (75 mg) depots were homogenized in Zurich medium (10 mM Tris-HCl, 1 mM EDTA, 150 mM NaCl, 1% Triton X-100, 1% sodium cholate, 1% SDS) with a cocktail of protease inhibitors. Samples were centrifuged (Eppendorf, Madrid, Spain) and supernatant was collected and frozen at –80 °C until further use. Samples of the homogenates and the pre-stained molecular weight markers (Precision, BioRad, Madrid, Spain) were separated by SDS electrophoresis and electro-transferred to polyvinylidene difluoride membranes (0.45 μM, Millipore, Spain). After 1h blocking in milk, the membranes were incubated overnight at 4°C with the primary antibodies against PGC-1α (1:1000; 92 kDa; Santa Cruz Biotechnology INC, Texas, EUA), PPARγ (1:1000; 53-57 kDa, Cell Signaling Technology, Massachusetts, EUA), HSL (1:1000; 83 KDa; Cell Signaling Technology, Massachusetts, EUA), pATGL (phospho S406) (1:1000; 55 KDa; Abcam, Cambridge, UK), pAMPKα (phospho Thr172) (1:1000; 60 kDa; Cell Signaling Technology, Massachusetts, EUA), TH (1:1000; 60 kDa; Abcam, Cambridge, UK), β2 receptors (1:200; 47 KDa; Alomone, Jerusalem, Israel) β3 receptors (1:200; 45KDa; Alomone, Jerusalem, Israel), D1R (1:200; 48 KDa; Abcam, Cambridge, UK), D2R (1:200; 49 KDa, Sigma-Aldrich, Madrid, Spain). The membranes were washed with Tris-buffered saline with Tween (TBST) (0.1%) and incubated with rabbit anti-goat (1:5000; Thermofisher Scientific, Massachusetts, EUA), goat anti-mouse (1:5000; Bio-Rad Laboratories, California, EUA) or goat anti-rabbit (1:5000; Bio-Rad Laboratories, California, EUA) in TBS and developed with enhanced chemiluminescence reagents (ClarityTM Western ECL substrate, Hercules, CA, USA). Intensity of the signals was detected in a Chemidoc Molecular Imager (Chemidoc; BioRad, Madrid, Spain) and quantified using Image Lab software (BioRad). The membranes were re-probed and tested for Calnexin (1:1000; 90 KDa; SICGEN, Cantanhede, Portugal) and/or β-actin (1:1000; 42 KDa; SICGEN, Cantanhede, Portugal) immunoreactivity to compare and normalize the expression of proteins with the amount of protein loaded.

### Histological and immunohistochemical evaluation of WAT and BAT

Visceral WAT and BAT depot were collected, dissected and immersed-fixed in PFA 4%. Samples were then embedded into paraffin (Sakura Finetek Europe B.V., Zoeterwoude, Netherlands) and longitudinal serial sections of 8 or 10 μm thick were obtained with a Microtome Microm HM200 (MICROM Laborgeräte GmbH, ZEISS Group, Walldorf, Germany).

For evaluation of adipocytes perimeter, after sectioning, the samples were transferred into slides and stained with hematoxylin and eosin, for staining of the nuclei, extracellular matrix and cytoplasm. Representative photographs were acquired using NDP.view2 software (Hamamatsu, Japan) in slides digitally scanned in the Hamamatsu NanoZoomerSQ (Hamamatsu, Japan). Adipocyte’s perimeter was visualized with software Fiji app for Image J (https://imagej.nih.gov/ij/).

UCP1 protein and mitochondrial density were evaluated by immunohistochemistry. Paraffin sections were deparaffinized and rehydrated followed by antigen retrieval and blocking, performed with a 5% bovine serum albumin solution for 60min. Sections were then incubated with primary antibody rabbit anti-UCP1 (1:1000 for WAT and BAT; Santa Cruz Biotechnology INC, Texas, EUA), overnight at 4°C. Next, sections were incubated with anti-rabbit secondary antibody Alexa 488 (1:4000 for WAT and 1:6000 for BAT; Termofisher Scientific, Massachusetts, EUA), Mitotracker^TM^ Red CMXRos [15nM] for WAT and [1nM] for BAT (Termofisher Scientific, Massachusetts, EUA) and DAPI (1ug/ml; Santa Cruz Biotechnology INC, Texas, EUA) for 90min. Finally, sections slides were mounted with Dako mounting medium (Agilent, California, EUA), visualized at the widefield Z2 Zeiss Microscope (ZEISS Group, Walldorf, Germany) and analyzed with software Fiji app for Image J (https://imagej.nih.gov/ij/).

### Evaluation of oxygen consumption rate

Three weeks after CSN resection, mice were sacrificed by cervical dislocation and WAT and BAT depots were collected for the evaluation of oxygen consumption rate (OCR) using the Seahorse XF24 (Seahorse Bioscience, North Billerica, MA). Tissue samples were placed in an XF24 Islet Capture Microplate (Seahorse Bioscience, North Billerica, MA) and once in position, were rinsed twice with Seahorse XF DMEM assay media (Seahorse Bioscience, North Billerica, MA) supplemented with 10mM glucose, 1mM pyruvate and 2mM L-glutamine. Finally, 575μl of assay media was added to all the samples and control wells. Before the measurement of the OCR, the microplate was incubated at 37°C without CO2 for 45 min. To evaluate adrenergic stimulation, OCR was measured after norepinephrine [15μM] (Sigma-Aldrich, Lisboa, Portugal) or dopamine [100nM] (Sigma-Aldrich, Lisboa, Portugal) application. OCR was calculated by plotting the O2 tension of the media as a function of time (pmol/min).

### *In vivo* tissue-specific glucose uptake evaluation

An intravenous glucose tolerance test (IVGTT) was performed in sham versus CSN - transected normal chow and HF animals. For that, the animals were fasted overnight and a bolus of 2-deoxy-D-[1,2-3H]-glucose (1mC/ml; specific activity: 20Ci/mmol; PerkinElmer, Madrid, Spain) mixed with glucose (100µCi/kg body weight; 0.5g/kg body weight) was administered in the tail vein. Blood samples were taken from the tail vein at regular intervals (0, 2, 5, 10, 15, 30 and 60 minutes).

To determine glucose-specific activity, 20µl plasma was deproteinized with 200µl ice-cold perchloric acid (0.4N), centrifuged and radioactivity was measured in a scintillation counter (Tri-Carb 2800TR, Perkin-Elmer, Madrid, Spain). At 60 minutes, animals were euthanized, and the tissues (white and brown adipose tissue depots) were rapidly excised. 2-deoxy-D-[3H]glucose incorporation was investigated in 50-200mg tissue homogenized in 1ml ice cold perchloric acid (0.4 N)(6). The samples were centrifuged and the radioactivity in the supernatant was measured in a scintillation counter.

### Light-sheet based fluorescent microscopy

After perfusion, the excised BAT and WAT were washed with PBS and clarified using the iDisco+ method (https://idisco.info/). Briefly, samples were dehydrated at room temperature in successive washes of 20% Metanol (MetOH) for 1h, 40% MetOH for 1h, 60% MetOH for 1h, 80% MetOH for 1h, 100% MetOH for 1h and 100% MetOH overnight. After, the samples were incubated in a solution of 33% MetOH and 66% Di-ChloroMethan (DCM, Sigma) overnight and washed twice with 100% methanol for 1h. Samples were then bleached in chilled fresh 5% H_2_0_2_ in methanol overnight, at 4°C, before being rehydrate with methanol/H_2_O series (80%, 60%, 40%, 20% and PBS, 1 h each at room temperature). Samples were them immunolabeled for 24 h with anti-TH (AB152, Merck, France) after overnight permeabilization at 37°C and overnight blocking with 1,7% TritonX-100, 6% donkey serum and 10% DMSO in PBS. After 3 washes in PBS/ 0.2%Tween-20, samples were incubated with the secondary antibody donkey anti-goat (Jackson Immunoresearch, Cambridgeshire, UK) for 24h. Samples were then dehydrated at room temperature in successive baths as described above and then incubated in a solution of 33% MetOH and 66% Di-ChloroMethan (DCM, Sigma) for 3h at RT, then in 100% DCM twice for 15 min twice and transferred overnight into the clearing medium 100% DiBenzylEther 98% (DBE, Sigma).

Imaging was performed using a home-made light-sheet ultramacroscope. The specimen was placed into a cubic cuvette filled with DBE placed on the Z-stage of the bench. It was illuminated with planar sheets of light, formed by cylinder lenses. The light coming from a multi-wavelength (561 nm) laser bench (LBX-4C, Oxxius) was coupled via two single mode optical fibers into the setup, allowing illumination from one or two sides. Two-sided illumination was used. The specimen was imaged from above with a MVX10 macroscope, through a PlanApo 2X/0.5 NA objective (Olympus) with an additional zoom of the macroscope of 1.6, which was oriented perpendicular to the 561nm light sheet.

Images were captured using a sCMOS Camera (Orca-Flash4.0) synchronized with the z-stage moving the sample through the light sheet. The ultramicroscope is managed by Micro-manager software and z-stacks of images were taken every 2 μm. The images stacks were fused using the alpha-blending method with a home-made ImageJ macro (Rasband, W.S., ImageJ, U. S. National Institutes of Health, Bethesda, Maryland, USA, http://imagej.nih.gov/ij/, 1997e2012).

### Free-floating **Δ**Fos immunohistochemistry in the paraventricular nucleus of the hypothalamus

Cryopreserved brains of sham vs CSN-transected NC and HF animals were cut in coronal sections (40μm) using a cryostat and collected for a 6-well plate with 4 sections/well in PBS. Coordinates used for sectioning were -2.16mm and - 1,8mm to evaluate different regions of the paraventricular nucleus (PVN) of the hypothalamus. Free-floating immunofluorescence was performed in all evaluated slices. Briefly, sections were transferred to a 24 wells plate (1 section/well) and gelatin was removed by incubating sections for 1h at 37°C. Sections were then permeabilized in 0.1M TBS, 0.3% Triton X-100 for 3×15 minutes in a shaker at room temperature and washed in TBS-T. Unspecific binding was blocked with 1% BSA, 0.3% Triton x-100, 3% normal goat serum in 0.1 M TBS, for 2h at 24°C. After, slices were incubated with primary antibody for delta-FosB (1:100; Cell signaling Technology, Massachusetts, EUA) overnight at 4°C. Negative control was performed by avoiding incubation with primary antibody and kept in blocking solution for this period. Solution was decanted and slices washed in TBS-T before being incubated with specific secondary antibody in a dilution of 1:2000 for 1h at 24°C in the dark (anti-rabbit Alexa Fluor 550 or anti-mouse Alexa Fluor 488; from Abcam, Cambridge, UK). Slices were washed in TBS in the dark and mounted with a coverslip and a drop of mounting medium Fluoroshield™ - Sigma-Aldrich. Each slide was immediately visualized in a confocal Zeiss LSM980 microscope and fluorescence images were used to measure the percentage of delta-FosB-immunoreactive cells per total number of cells.

### Statistical Analysis

Data were evaluated using GraphPad Prism Software, version 9 (GraphPad Software Inc., San Diego, CA, USA) and presented as mean values with SEM. The significance of the differences between the mean values was calculated by one- and two-way ANOVA with Bonferroni multiple comparison test. Differences were considered significant at *p* < 0.05.

## Data and code availability

Published data and supplemental information include all the data used to generate the figures in the paper. This paper does not report original code. Any additional information required to reanalyze the data reported in this paper is available upon request.

## Acknowledgments

We would like to acknowledge the Histology and Microscopy facilities from NOVA Medical School|Faculdade de Ciências Médicas, Universidade NOVA de Lisboa for their help in the preparation of adipose tissue slices for immunohistochemical analysis and to Oscar Rios-Rodriguez from the Departamento de Bioquímica, Biologia Molecular y Fisiologia, Universidad de Valladolid, Valladolid, Spain, for its help in the quantification of catecholamines by HPLC.

## Author contributions

The authors have contributed to the study as follows. Participated in research design, S.V.C., P.B.; conducted experiments, B.F.M., J.F.S, F.O.M., C.P. A.F.,

D.R., J.L., E.O., M.C., S.V.C.; performed collection and data analysis, B.F.M., J.F.S, F.O.M., C.P. A.F., D.R., J.L., E.O, G.M.M., P.M., J.P., A.R., S.V.C., P.B.; wrote or contributed to the original writing of the manuscript, B.F.M., J.F.S. S.V.C; contributed to the review & editing of the manuscript, B.F.M., J.F.S., P.B, S.V.C. All the authors have approved the final version of the manuscript and agree to be accountable for all aspects of the work in ensuring that questions related to the accuracy or integrity of any part of the work are appropriately investigated and resolved. All authors have read and agreed to the published version of the manuscript.

