## Supplemental table 1 and Figure 1 for "Ablation of carotid body activity reverses diabesity by improving white and brown adipose tissue sympathetic innervation and metabolism"

### Supplemental data

**Table S1** - Effect of CSN resection on insulin sensitivity, glucose metabolism and lipid profile on obese dysmetabolic rats and mice

| Rats | | | | | |
|---|---|---|---|---|---|
| | | | **Before diet** | **Before surgery** | **9 weeks after surgery** |
| **Caloric intake (Kcal/day/kg)** | NC | Sham | - | 197.63±9.68 | 218.77±30.21 |
| | | Denervated | - | 215.15±7.14 | 217.22±19.91 |
| | HF | Sham | - | 280.08±29.76$^{*}$ | 301.63±17.18$^{*}$ |
| | | Denervated | - | 274.63±25.69$^{*}$ | 248.45±21.34 |
| **Glycemia (mg/dl)** | NC | Sham | 95.22±5.27 | 75.11±3.48 | 82.90±3.60 |
| | | Denervated | 85.66±3.77 | 84.44±4.38 | 85.22±3.71 |
| | HF | Sham | 78.22±3.16 | 95.11±5.06$^{*}$ | 92.90±2.20$^{\S\S}$ |
| | | Denervated | 81.56±2.12 | 98.78±4.10$^{**}$ | 81.5±2.46$^{\#\#}$ |
| **Insulin sensitivity Kitt (%glucose/min)** | NC | Sham | 4.43±0.24 | 4.24±0.24 | 4.78±0.31 |
| | | Denervated | 5.03±0.33 | 4.50±0.25 | 4.85±0.31 |
| | HF | Sham | 4.1±0.21 | 1.76±0.33$^{***, \#\#\#\#}$ | 1.84±0.23$^{\S\S\S, \#\#\#\#}$ |
| | | Denervated | 4.7±0.35 | 1.92±0.38$^{***}$ | 4.24±0.20$^{\#\#\#}$ |
| **Glucose Tolerance AUC OGTT (mg/dl*min)** | NC | Sham | 20747.55±692.56 | 18439.55±810.08 | 19966.89±449.27 |
| | | Denervated | 19286±498.99 | 18160.22±770.48 | 20813.45±662.14 |
| | HF | Sham | 21028.67±759.24 | 25359.45±825.60$^{***, \#\#}$ | 24504.63±355.01$^{\S\S}$ |
| | | Denervated | 20116.11±746.74 | 25415.33±689.33$^{***, \#\#\#\#}$ | 23264.29±433.77$^{\#}$ |
| **Insulin (pmol/l)** | NC | Sham | 5.24±1.34 | 170.39±9.20 | 240.55±16.00 |
| | | Denervated | 19.43±2.06 | 146.68±15.43 | 227.78±17.16 |
| | HF | Sham | 22.98±4.06 | 415.58±35.34$^{****, \#\#\#\#}$ | 584.21±45.95$^{\S\S\S\S}$,$^{\#\#\#\#}$ |
| | | Denervated | 25.87±1.79 | 506.94.77±22.48$^{****, \#\#\#\#}$ | 420.92±21.69$^{\S\S\S\S,\#\#\#\#}$ |
| **C-peptide (nmol/l)** | NC | Sham | 0.33±0.09 | 0.68±0.16 | 0.64±0.08 |
| | | Denervated | 0.45±0.12 | 0.72±0.08 | 0.76±0.06 |
| | HF | Sham | 0.33±0.06 | 1.34±0.07$^{****, \#\#\#\#}$ | 1.83±0.17$^{\S\S\S\S}$,$^{\#\#\#}$ |
| | | Denervated | 0.37±0.10 | 1.51±0.16$^{****, \#\#\#\#}$ | 1.34±0.15$^{\#}$ |
| **Cholesterol (mg/dl)** | NC | Sham | - | - | 61.20±4.47 |
| | | Denervated | - | - | 63.84±1.54 |
| | HF | Sham | - | - | 95.65±4.56$^{****}$ |
| | | Denervated | - | - | 77.33±4.14$^{\#}$ |
| **Triglycerides (mg/dl)** | NC | Sham | - | - | 90.65±7.02 |
| | | Denervated | - | - | 91.96±8.21 |
| | HF | Sham | - | - | 147.05±24.07$^{*}$ |
| | | Denervated | - | - | 108.12±11.06 |
| **c-LDL (mg/dl)** | NC | Sham | - | - | 4.08±0.31 |
| | | Denervated | - | - | 4.32±0.17 |
| | HF | Sham | - | - | 7.97±1.06$^{***}$ |
| | | Denervated | - | - | 5.87±0.54 |
| **NEFA (mg/dl)** | NC | Sham | - | - | 1.06±0.08 |
| | | Denervated | - | - | 1.10±0.07 |
| | HF | Sham | - | - | 1.19±0.07 |

| | | | | | |
|---|---|---|---|---|---|
| | | Denervated | - | - | 0.70±0.18# |
| **Mice** | | | | | |
| | | | **Before diet** | **Before surgery** | **9 weeks after surgery** |
| **Caloric intake (Kcal/day/kg)** | NC | Sham | - | 353.00±12.79 | 365.70±11.90 |
| | | Denervated | - | 363.20±14.78 | 388.20±13.73 |
| | HF | Sham | - | 526.30±46.82** | 431.40±57.42 |
| | | Denervated | - | 490.50±43.78* | 457.70±40.35 |
| **Glycemia (mg/dl)** | NC | Sham | - | 80.60±2.42 | 86.40±2.79 |
| | | Denervated | - | 79.00±3.86 | 80.33±1.54 |
| | HF | Sham | - | 124.50±5.38*** | 144.75±10.06 |
| | | Denervated | - | 112.40±7.75** | 94.75±11.84 |
| **Insulin sensitivity AUC ITT (mg/dl*min)** | NC | Sham | - | 10641.20±864.94 | 9276.60±488.69 |
| | | Denervated | - | 11135.83±584.78 | 10477.50±1247.50 |
| | HF | Sham | - | 21645.75±1684.90*** | 23392.00±695.97 |
| | | Denervated | - | 22336.20±2069.03**** | 15251.80±1997.12#, § |
| **Glucose Tolerance AUC OGTT (mg/dl*min)** | NC | Sham | - | 23194.80±678.63 | 19657.60±971.99 |
| | | Denervated | - | 20851.33±2783.03 | 18667.83±535.42 |
| | HF | Sham | - | 32773.25±1155.82*** | 30596.35±1111.42 |
| | | Denervated | - | 33064.60±2783.03** | 26894.00±1191.86 |

Values represent means ± SEM; Two-Way ANOVA with Bonferroni multicomparison test. *$p<0.05$, **$p<0.01$, ***$p<0.001$ and ****$p<0.0001$ comparing normal chow animals and HF values; #$p<0.05$, ##$p<0.01$, ###$p<0.001$ and ####$p<0.0001$ comparing values without and with CSN resection; §$p<0.05$, §§$p<0.01$, §§§$p<0.001$, §§§§$p<0.0001$ comparing values 9 weeks after CSN with values 9 weeks after sham. AUC- area under the curve; c-LDL – low density lipoproteins cholesterol; HF – high fat; Kitt – constant of the insulin tolerance test; OGTT – oral glucose tolerance test; NEFA- non-esterified free fatty acids.

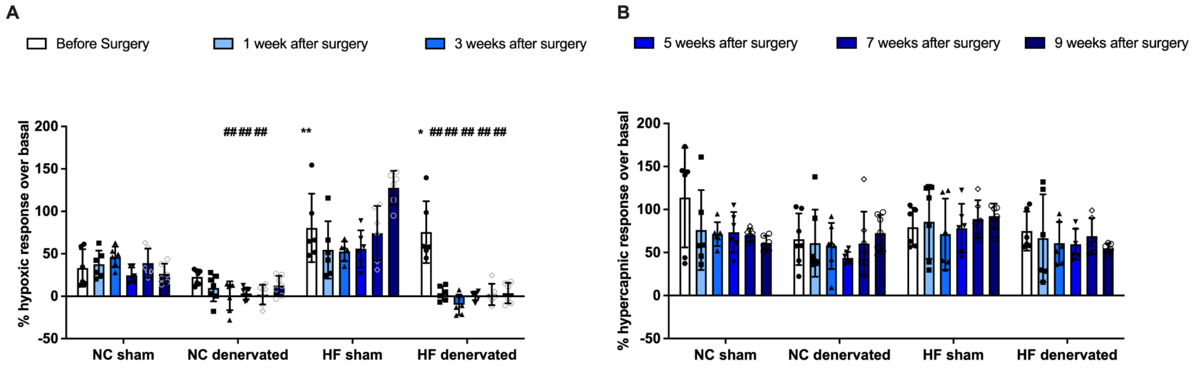

**Figure S1 – Carotid Sinus Nerve (CSN) resection decreases respiratory responses to hypoxia:** A) respiratory responses to hypoxia; B) respiratory responses to hypercapnia; Protocol used consisted in animal acclimatization during 30 min followed by 10 min of normoxia (20% O2 balanced N2), 10 min of hypoxia (10% O2 balanced N2), 10 min of normoxia, 10 min of hypercapnia (20% O2 + 5% CO2 balanced N2), and finally to 10 min of normoxia. Bars represent means ± SEM (n=9-15); Two-Way ANOVA with Bonferroni multicomparison test. *p<0.05, comparing NC and HF animals without CSN resection; $^{\#}$p<0.05 comparing groups before and after CSN resection.
